# No evidence for differences in contrast of the grey-white matter boundary in Autism Spectrum Disorders: An open replication

**DOI:** 10.1101/750117

**Authors:** Nicolas Traut, Marion Fouquet, Richard Delorme, Thomas Bourgeron, Anita Beggiato, Roberto Toro

## Abstract

The contrast of the interface between the cortical grey matter and the white matter is emerging as an important neuroimaging biomarker for several brain disorders. Differences in grey to white matter contrast could be related to abnormalities in neuronal migration or in intra-cortical myelination, and are an appealing biomarker for ASD. Two previous studies have reported differences in contrast between patients with autism spectrum disorder and non-autistic controls.

We aimed at replicating this finding using open data from the ABIDE initiative, phases 1 and 2, gathering data from 2,148 subjects from 26 different centres and on 764 individuals from the EU-AIMS project (6 different centres). We used multiple linear regression to study the effect of the diagnosis of ASD on contrast, and 3 different strategies for controlling for multiple comparisons. We did not find statistically significant differences in the EU-AIMS dataset, and those that we found in the ABIDE dataset were due to a single centre. All the code necessary to replicate our analyses has been made available open source: https://github.com/neuroanatomy/GWPC.

## Introduction

Autism Spectrum Disorder (ASD) is characterised by persistent deficits in social communication and social interaction, and by restricted, repetitive patterns of behaviour, interests, or activities, that can be associated with hyper or hypo sensoriality. According to data released by the United States Center for Disease Control, the autism prevalence in the United States of America is 1 in 59 children (Baio 2018). This high prevalence and the significant disability that results from these symptoms make ASD a top health research priority.

Structural magnetic resonance imaging (MRI) is a powerful tool to explore the neuronal correlates of ASD. Structural MRI data can be acquired with sedation, which allows for the exploration of individuals with diverse intellectual abilities. A large number of reports exist of structural differences in the neuroanatomy of individuals with ASD compared with controls. In particular, many studies have reported an increased cortical thickness and grey matter (GM) volume, and decreased white matter (WM) volume in subjects with ASD. For example, in a longitudinal study Lange et al. (2015) reported an initial increase in GM volume in childhood, followed by a decrease of this volume, with a crossing of the control subjects curve between 10 and 15 years. The GM volume of many structures continued to decline in adults with ASD (Lange et al. 2015). In addition, other results evoke a notable bilateral regional decrease in GM volume in the amygdalo-hippocampal complexes and in the precuneus (Via et al. 2011). But the definition of the GM and WM volumes in MRI is necessarily conditioned by our ability to place the grey-white boundary, itself derived from the contrast between grey and white matter tissue signal intensities.

The cortical thickness abnormalities observed could therefore reflect a difference in grey-to-white matter percent contrast (GWPC) between patients and controls. This difference could in turn reflect differences in the cellular composition of the grey matter, for example neuronal migration abnormalities, or differences in intra-cortical myelination. Several studies have suggested that ASD may stem from abnormalities in brain connectivity, and such differences in myelination could be a proxy for connectivity disorders. Grey-to-white matter contrast appears thus as an appealing biomarker for ASD.

To date, two studies have looked at differences in grey-to-white matter contrast. Andrews et al. (2017) analysed the contrast of the grey-white matter boundary in a group of 98 ASD patients and 98 matched controls. They reported a significant decrease in contrast at different cortical levels in ASD patients. Olafson et al. (2021) analysed the slope of grey level at the white matter boundary in a group of 1136 individuals. They reported a significant increase in contrast in ASD patients in the bilateral superior temporal gyrus and left inferior frontal gyrus.

We aimed at replicating the reports of contrast abnormalities using a large open sample: the *Autism Brain Imaging Data Exchange* 1 and 2 projects (ABIDE 1 and ABIDE 2), which includes data from 2,148 individuals (1,019 individuals with ASD and 1,129 controls from the general population). These data were made available in March 2017 (Di Martino et al. 2014, 2017). The ABIDE project data has proven useful in the past to attempt open replications of previous findings (Traut et al. 2017; Haar et al. 2016; Lefebvre et al. 2015; Heiss et al. 2015). The data in ABIDE 1 and 2 comes from 26 international sites, sharing previously collected T1-weighted anatomical MRI data, plus phenotypic information for each participant. The acquisition protocols at each site have been developed independently, and some researchers have criticised the results obtained using ABIDE data for this lack of harmonisation. To address this issue, we performed an additional replication, using exactly the same methods, in 764 subjects from the EU-AIMS project, where many efforts have been made to harmonise data collection. Our main objective was to study the presence of differences in GWPC between individuals with a diagnosis of ASD and non-ASD controls. Additionally, we studied the impact of age, sex, intellectual quotient (IQ), and the relationship between GWPC and cortical thickness.

## Materials and Methods

### Study participants

We analysed all individuals from the ABIDE 1 and 2 projects, N=2,148. There were 1,019 individuals with ASD (diagnosis confirmed on standardised instruments) and 1,129 controls with a neurotypical development (http://fcon_1000.projects.nitrc.org/indi/abide). The number of MRIs was larger than the number of participants (2,214 MRIs for 2148 subjects) because some individuals had undergone several MRIs. We replicated our analysis using time point 1 of the EU-AIMS dataset, containing 764 participants (453 with ASD and 311 controls) from 6 different centres (https://www.eu-aims.eu). Access to EU-AIMS data can be requested online.

### Cortical surface reconstruction

We preprocessed all available structural MRI data using a homogeneous pipeline. Data was acquired on 3 Tesla scanners, with the exception of 1 centre from ABIDE where it was acquired in a 1.5 Tesla scanner. We used the surface reconstruction pipeline implemented in FreeSurfer v6.0 (Segonne et al. 2004; Dale et al. 1999; Jovicich et al. 2006). In brief, after intensity normalisation and brain extraction, the tissue was segmented into grey and white matter, and an atlas-driven approach was used to label different brain regions. Topologically spherical meshes were then built which reconstructed the boundary between grey and white matter (i.e., the “white matter” surface), and between the grey matter and the cerebro-spinal fluid (i.e., the pial surface). The placement of the grey-white matter boundary is based on an algorithm that aims at detecting the largest shift in signal intensity. By using intensity gradients across tissue classes the boundary placement is not reliant solely on absolute signal intensity and allows for subvoxel resolution in the placement of the grey-white matter surface (Fischl and Dale 2000).

### Grey-to-white matter percent contrast (GWPC)

Once a grey-white matter surface was obtained, the GWPC was calculated from a measure of the WM signal intensity at 1.0 mm into the white matter from the grey-white interface, and at intervals of 10% up to 60% of the distance from the white matter to the pial surface, thus yielding a set of 7 grey matter intensities (i.e., 0 – 60%). The outer section (i.e., 70 – 100%) of the cortical sheet was not sampled to ensure that sampling was performed within the cortical grey matter alone, and not confounded by voxels including cerebrospinal fluid (Fig. 1). The GWPC is the calculation of a ratio between GM and WM intensities:

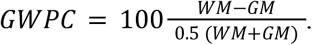

**Figure 1.**
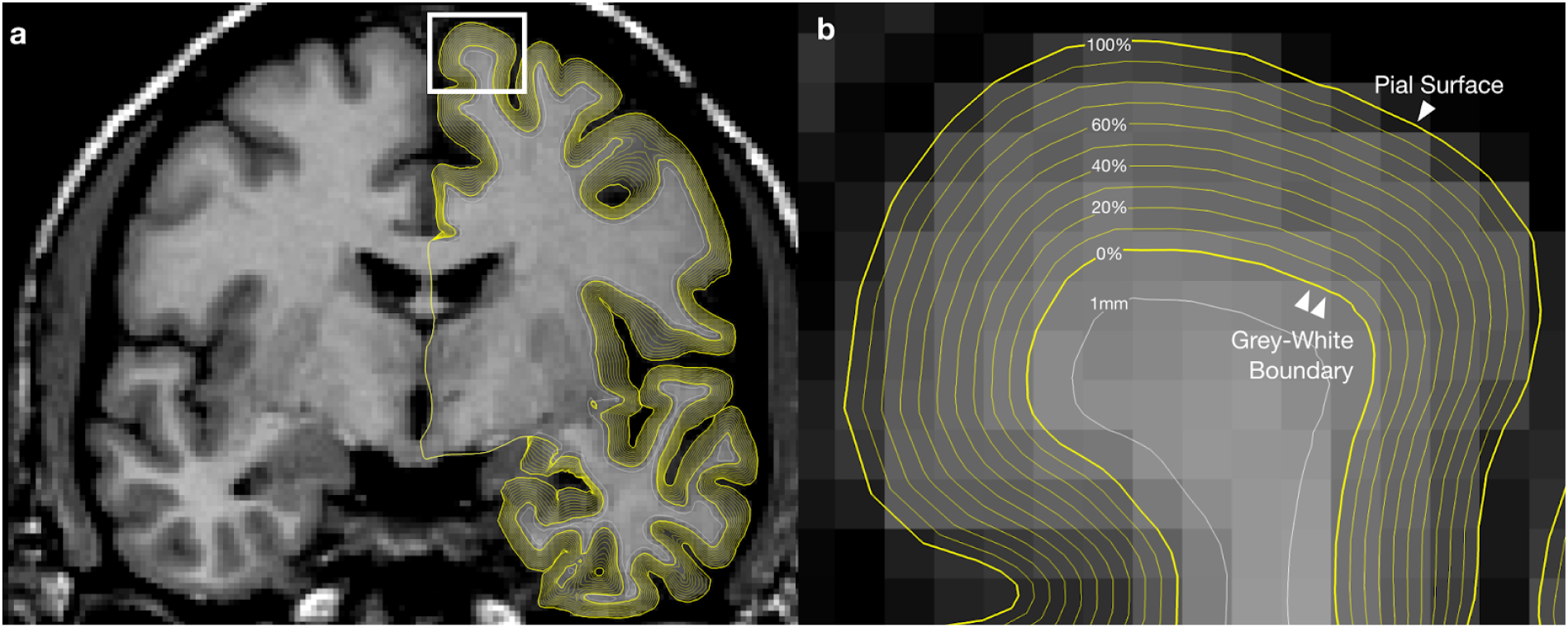
Selection procedure for white matter intensity and grey matter intensities. **a**. Hemisphere showing the different surfaces across the grey matter and the surface inside the white matter where grey-level values were sampled. **b**. Enlargement of the region framed in white in Fig. 1a: The 0% surface represents the grey-white boundary surface, the 100% represents the pial surface. The white matter signal intensity was sampled in a surface located 1mm below the grey-white boundary.

### Quality control

We visually controlled for the quality of the MRI data, segmentations, and surface reconstructions. First, we excluded MRIs for which the FreeSurfer automatic segmentation had failed (most often due to poor raw data quality). We then controlled the automatic segmentation performed by FreeSurfer. Whenever a subject had more than 1 MRI, we selected the one with the best data quality and segmentation.

We used QCApp to control the quality of the segmentations produced by FreeSurfer (available at https://github.com/neuroanatomy/QCApp). This application allowed us to quickly browse through the complete list of individuals, providing sagittal, coronal and axial views. An example of the display is shown in Fig. 2a. Each image is a full stereotaxic viewer, allowing to investigate further slices in case of doubts about a segmentation. A panel on the side provides z-scores of the volume of different structures segmented relative to the averages and standard deviations of the complete population.

**Figure 2.**
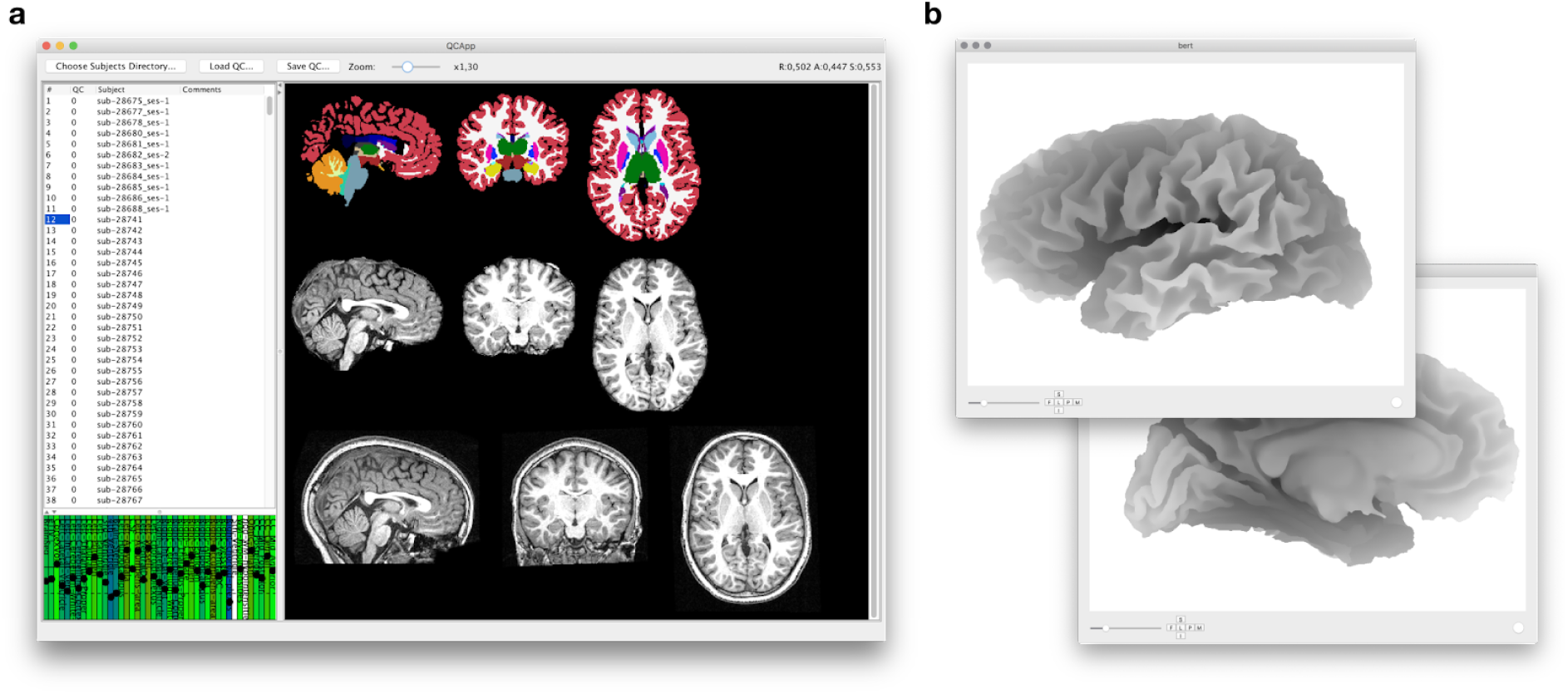
a : Appearance of Good Quality Segmentation by FreeSurfer v6.0.0. Lower line: MRI before any treatment. Intermediate line: MRI after removal of signals corresponding to the scalp. Upper line: MRI after automatic segmentation by FreeSurfer. Left column: Sagittal view. Central column: coronal view. Right column: axial view. **b. Aspect of a grey-white interface reconstruction**. Lateral view (left) and medial view (right) of a left hemisphere from a good quality MRI.

We used SurfaceRAMON to visually control the quality of the pial surface reconstruction and the grey-white matter surface reconstruction (available at https://github.com/neuroanatomy/SurfaceRAMON). As before, this application provides a list of all individuals in the sample, and allows us to quickly browse through them. The images are full 3D objects which can be rotated to inspect specific regions. We controlled the quality of the segmentation by inspecting the lateral and medial views of the left and right hemispheres, and inspected different angles in case of doubt (Fig. 2b).

For the ABIDE analysis, after controlling for data quality we excluded individuals missing some of the data required for our statistical analyses: diagnosis, sex, age, or scanning centre. Total IQ was not available for 111 subjects. For 40 subjects among them we imputed total IQ from verbal and performance IQ using a linear regression model: totalIQ ∼ verbalIQ + performanceIQ. The regression of total IQ on verbal and performance IQ had an R^2^ = 0.95. The remaining 71 individuals were excluded from further analyses. Finally, we excluded 53 individuals with ASD from centres where no controls were available. The final sample included 1,475 individuals gathering 636 with ASD and 839 with neurotypical development. The inclusion/exclusion flow is described in Supplemental Fig. 1. Additional details on the number of subjects per site and the presence of comorbidities are available in Supplemental Tables 1 and 2.

For the EU-AIMS analysis, we trained a prediction model based on the FreeSurfer outputs on the quality control we did on the ABIDE dataset. Based on the segmentation quality prediction, we excluded 138 participants because of poor segmentation quality (100 ASD patients and 38 controls). We excluded 3 additional subjects because their full-scale IQ was missing. The final sample included 586 individuals (323 ASD, 263 controls). Additional details on the phenotype and number of subjects per site in the EU-AIMS sample are available in Supplemental Tables 3 and 4)

### Statistical analyses

To compute GWPC we first smoothed the MRI data using a filter with a full-width at half-maximum of 10 mm. We fit a linear regression model to predict GWPC measurements from diagnostic group (ASD or control), sex, scanning centre, age and IQ. We computed p-values using 2-tailed tests, and the statistical significance level was set to alpha = 0.05. We analysed the two hemispheres separately, which was taken into account using a Bonferroni correction. We accounted for multiple testing across brain vertices using Random Field Theory (RFT) as in the original article by Andrews et al. (2017). We additionally used Monte Carlo simulations and False Discovery Rate (Benjamini and Hochberg 1995). RFT analyses were performed using the Matlab/Octave SurfStat Toolbox (Worsley et al. 1999, http://www.math.mcgill.ca/keith/surfstat). For multiple comparison correction based on the RFT and Monte Carlo methods, clusters were formed for regions with a p-value <10^−2^. The significance of each cluster was evaluated based on the estimated number of independent resolution elements (resels) within the cluster for RFT, and the area of the cluster for Monte Carlo simulations. We preferentially looked at RFT analyses, and we used the Monte Carlo technique for analyses that were not feasible using the SurfStat Toolbox (for example, when we included cortical thickness as a covariate of the linear model). The results of RFT and Monte Carlo simulations were very similar. The analyses using false discovery rate (FDR) detected less significant findings.

Our first linear model included an interaction term between diagnosis and sex:

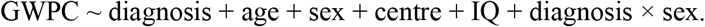

The interaction term was not significant in any of the analyses performed (after correction for multiple testing). We then preferred the analysis on a second linear model that did not include the interaction term, to avoid unnecessary loss of statistical power. The model we finally used was:

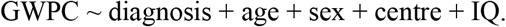

As a supplemental analysis, we tried to control for subject motion by including an estimate of this from resting state functional MRI as an additional covariate. We used the tool MCFLIRT (Jenkinson et al. 2002) to compute the root mean square deviation between consecutive volumes as a proxy measure of motion. The resulting model was as follows:

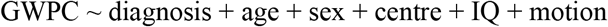

Vertex-based analyses were performed using FreeSurfer, and the results visualised using Freeview.

## Results

### Characteristics of the samples

There was no significant difference in age between the ASD and control groups (t=1.5918, p=0.11). There were proportionally less females in the ASD group than in the control group (24% vs 14%, *χ*_2_ (df=1)=25.0, p=5.7⋅10^−7^), as well as a significant difference in total IQ (lower in the ASD group, t=-10.219, p <2.2⋅10^−16^) (Table 1). Among the patients, only 7 had IQ <70. The rest of the analyses include total IQ as a covariate.

**Table 1.**
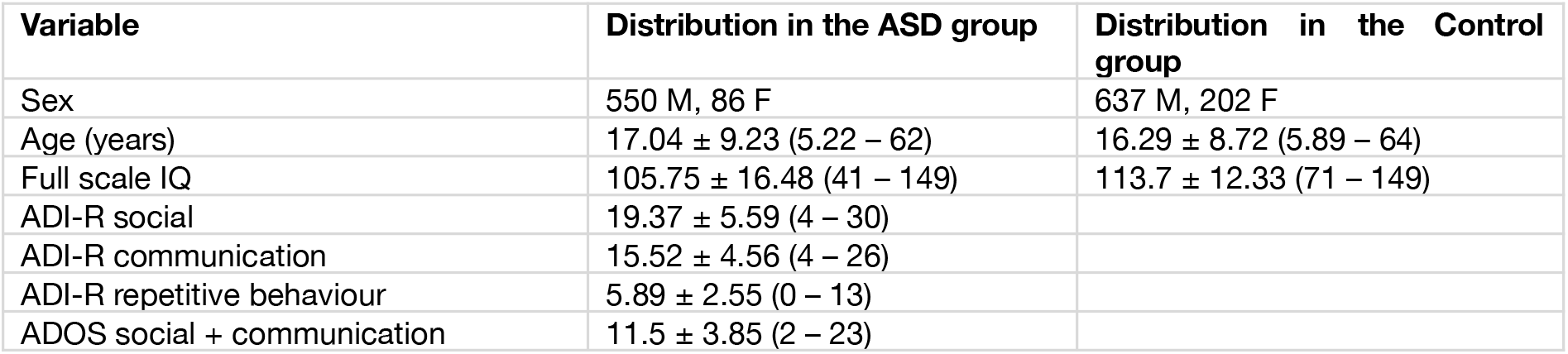
Characteristics of the ABIDE study sample. Results expressed as mean ± standard deviation (minimum, maximum). ADI-R: Autism Diagnostic Interview-Revised, ADOS: Autism Diagnostic Observation Schedule. ADI-R scores are missing for 208 ASD subjects and ADOS score is missing for 196 ASD subjects.

We included 15 centres from ABIDE 1, and 6 additional centres for Abide 2 (8 centres participated both in ABIDE 1 and 2). We further split the centre covariate when different MRI scanner acquisition protocols were used by the same centre, which was the case for OLIN, TRINITY, UCLA and KKI. Our quality control showed a subgroup of one KKI centre with very different MRI intensities: we added a specific centre level to tag this subgroup to ensure a certain degree of signal homogeneity within each group. In total, we defined 26 different levels of the centre covariate (see Supplemental Table 1). We excluded 521 subjects during quality control, i.e., 26% of the sample. Statistically significantly more subjects with ASD were excluded, 31% of the individuals with ASD versus 22% of the controls (*X*_2_(df=1)=21.325, p-value = 3.877⋅10^−6^). The subjects excluded were also significantly younger than those included: a median age of 12.25 years among the excluded versus 13.80 years in the included sample (Wilcoxon rank-sum test, p-value = 3.38⋅10^−11^). There was no difference in sex distribution between the excluded and included sample (26.2% of males were excluded versus 25.6% of females, *X*_2_(df=1)=0.040, p-value = 0.8409). The final ABIDE sample included 1475 subjects. Their characteristics are provided in Table 1.

**Table 2.**
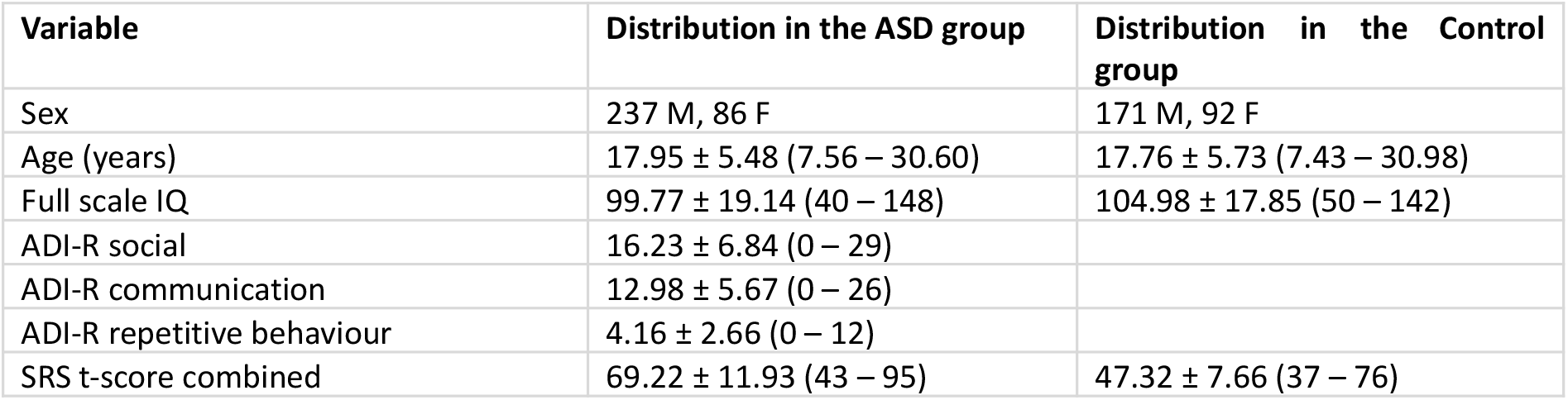
Characteristics of the study sample for the EU-AIMS dataset. Results expressed as mean ± standard deviation (minimum, maximum). ADI-R: Autism Diagnostic Interview-Revised, SRS: Social Responsiveness Scale. ADI-R scores are missing for 16 ASD subjects, SRS t-score is missing for 162 ASD subjects and 49 typical controls.

We included 586 subjects from the EU-AIMS project. Subjects were scanned in 6 different centres which were included as covariates. We excluded 138 individuals during quality control, 12.4% of the sample. The number of subjects with ASD excluded was statistically significantly higher (p-value < 0.00016). The subjects excluded were significantly younger (Wilcoxon rank-sum test, p-value = 2.3e-15). There was no significant difference in sex distribution between the excluded and included sample (12.4% of males were excluded versus 12.3% of females, *X*_2_(df=1)=2.7e-30, p-value = 1). The characteristics of the final EU-AIMS sample are provided in Table 2.

### No evidence for diagnosis group differences in GWPC

Our analysis of data from all 26 ABIDE centres did show clusters of statistically significant differences in GWPC between groups after correction with Monte Carlo simulations and RFT. However, after looking at the impact of each site on the effect, we realised that all the effect was due to a single site: NYU. Repeating the analysis after removal of NYU did not produce any statistically significant effect of diagnosis on GWPC. The non-thresholded maps of GWPC are shown in Fig. 3 (maps of the results obtained while including NYU are provided as Supplemental Fig. 2). Our analysis of the EU-AIMS dataset confirmed the absence of significant difference in GWPC due to ASD diagnosis (Supplemental Fig. 3).

**Figure 3.**
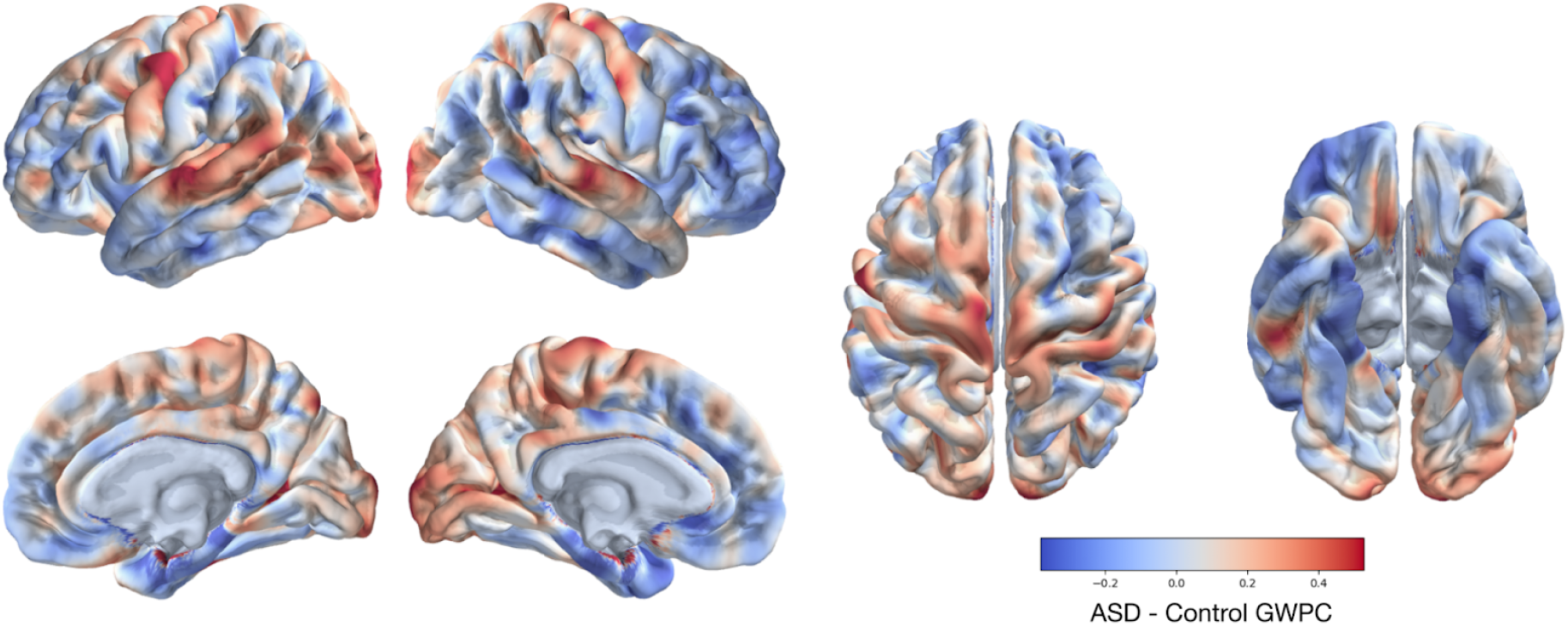
Non-thresholded map of the GWPC difference in ASD patients compared with control subjects. GWPC measured at 30% GM depth. There are no statistically significant differences.

### No evidence for diagnosis group differences in cortical thickness

The analysis of cortical thickness group differences including all 26 centres showed initially a set of statistically significant regions which overlapped with those of the GWPC analysis, only when correcting with RFT. Again, excluding the NYU site removed all statistically significant differences. Non-thresholded maps of differences in cortical thickness are shown in Fig. 4 (the results of the cortical thickness analysis including NYU are provided as Supplemental Fig. 4). The result was confirmed – no statistically significant difference in cortical thickness related to diagnosis group – when repeating the analysis using the EU-AIMS dataset (Supplemental Fig. 5).

**Figure 4.**
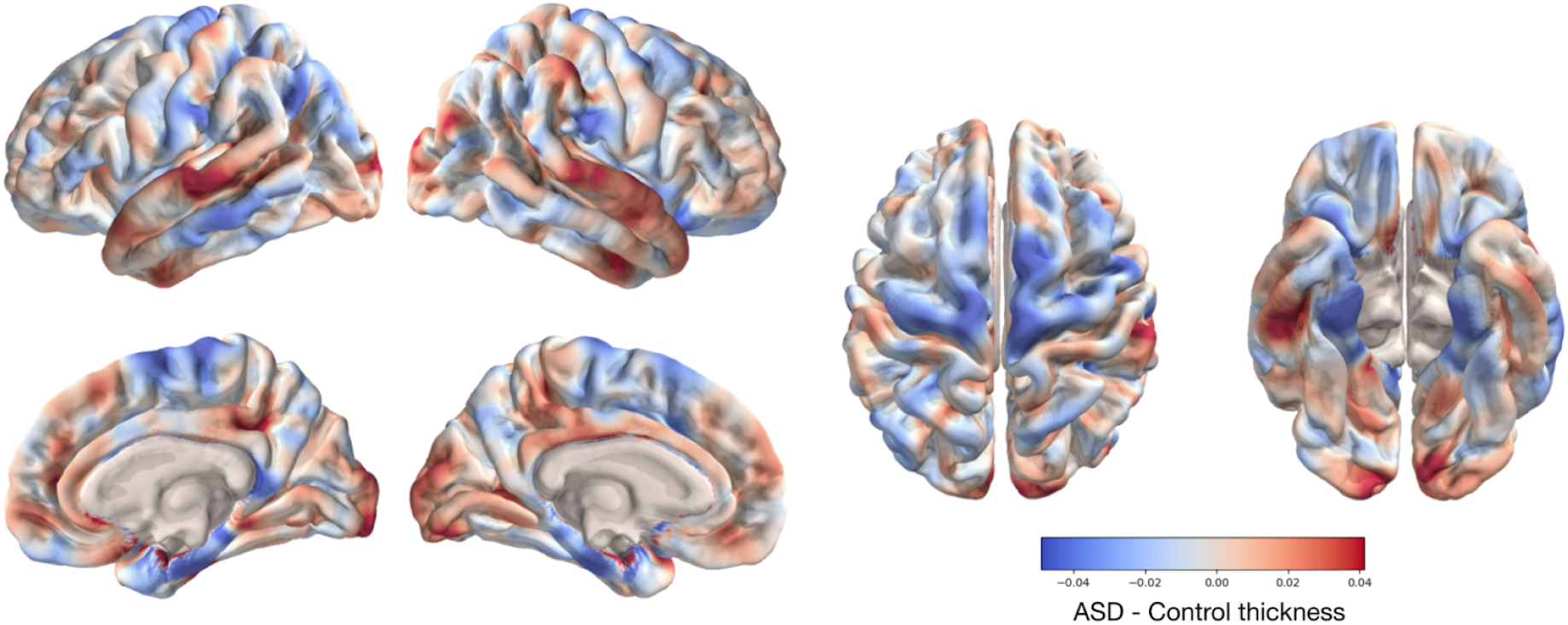
Non-thresholded maps of cortical thickness differences between ASD and control subjects. There are no statistically significant differences.

### Grey-to-white percent contrast is decreased in females compared with males

We observed several regions with decreased GWPC among females compared with males (Fig. 5, see Supplemental Table 5 for details of clusters with statistically significant decrease in GWPC among females), an effect that had already been reported (Salat et al. 2009; Andrews et al. 2017). The sex effect was robust to inclusion of cortical thickness as a covariate in the linear model (Supplemental Fig. 6). Most of the regions with significant differences revealed an increased cortical thickness in females compared with males, but they generally did not overlap with the regions with a reduced GWPC (Fig. 5). This result did not depend on the inclusion or exclusion of the NYU site, and was replicated in the EU-AIMS dataset (Supplemental Fig. 7).

**Figure 5.**
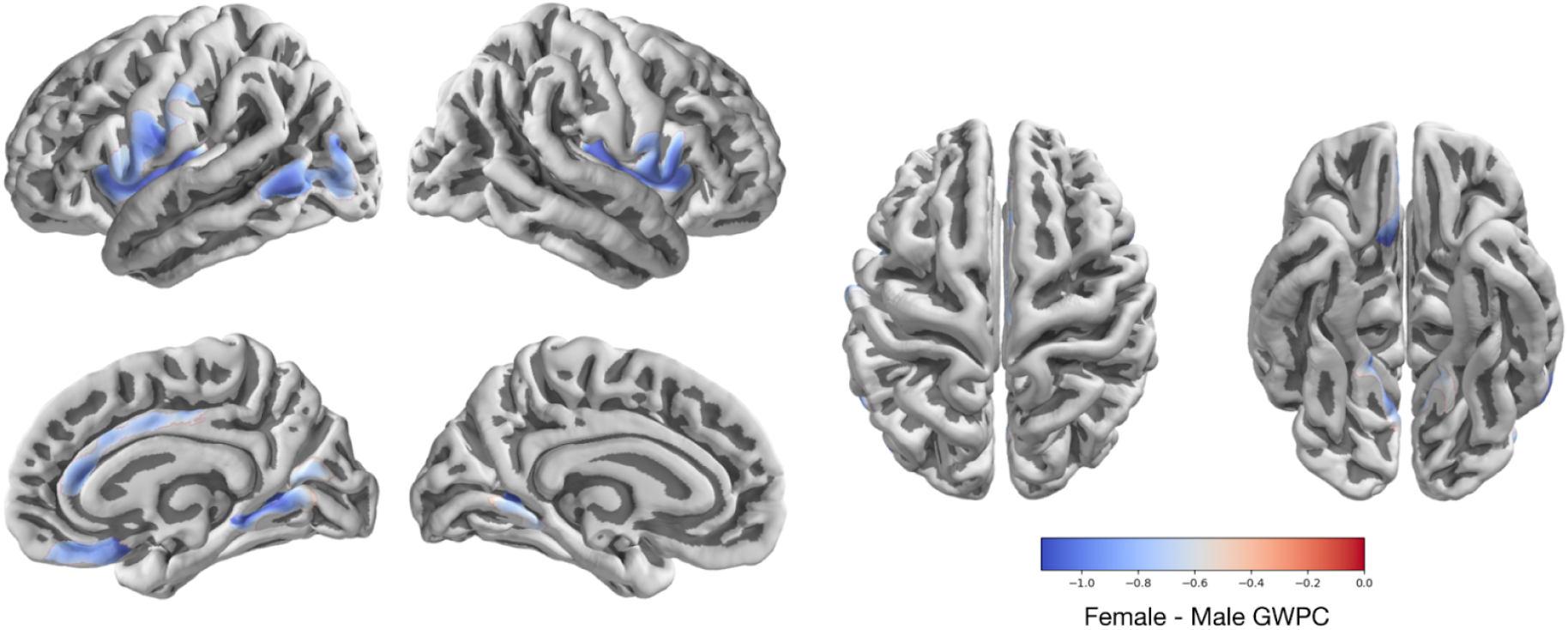
Significant decrease in GWPC in females compared with males. (statistically significant results at 5% after correction for multiple comparisons). Colour represents contrast, with negative values indicating Females<Males.

### Grey-to-white percent contrast decreases with age

We observed an overall decrease of GWPC with age (Fig. 6), consistent with results from the literature (Vidal-Piñeiro et al. 2016). The decrease in GWPC with age was particularly pronounced in the primary motor cortex. This could be potentially related to a continuation of intra-cortical myelination with age in our sample including young individuals. Cortical thickness also decreased with age over the entire cortex, with the exception of the insula and temporal areas where we observed an increase. Our results for a GWPC decrease remained significant after inclusion of cortical thickness as a covariate (see Supplemental Fig. 8 and Supplemental Table 6 for details of clusters p-values table). This result did not depend on the inclusion or exclusion of the NYU site, and was replicated in the EU-AIMS dataset (Supplemental Fig. 9).

**Figure 6.**
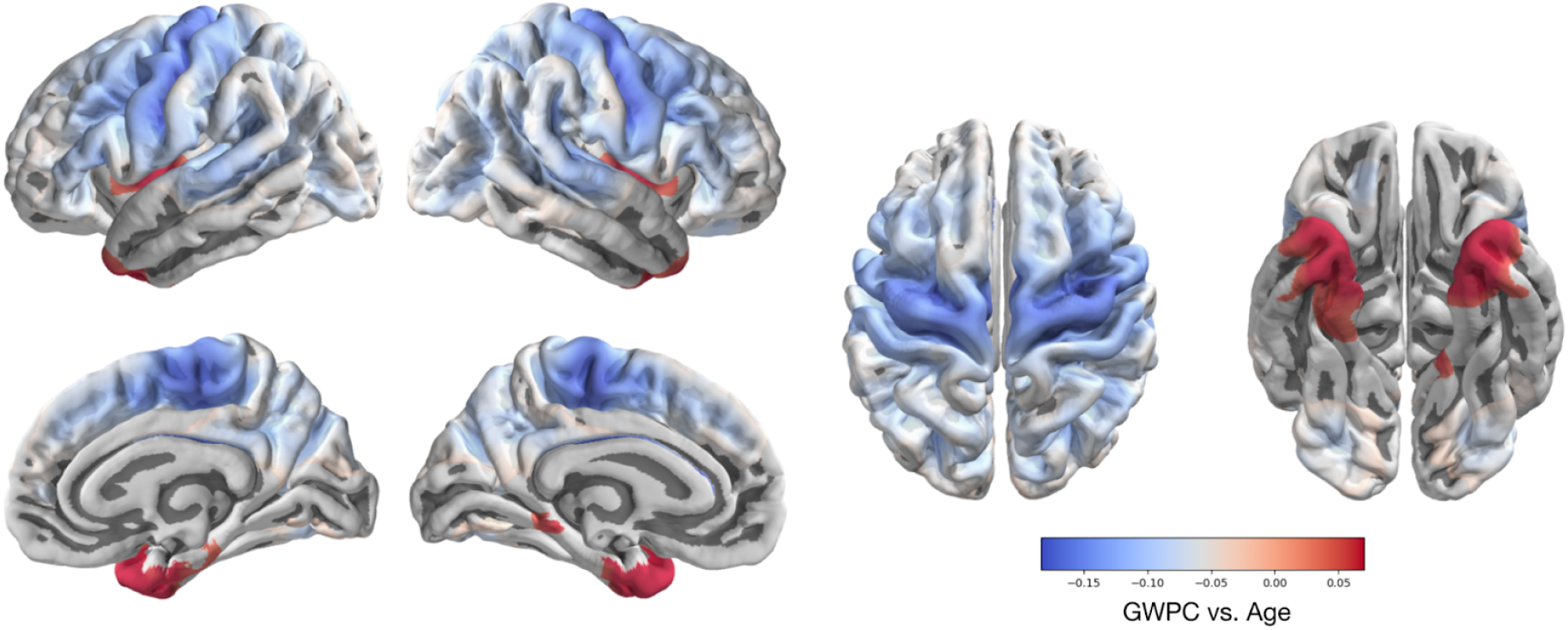
Significant changes in GWPC with age. GWPC shows an overall decrease with age, with the exception of regions within the insula and the temporal poles. Only significant results corrected for multiple comparisons are shown. Colours represent contrast, with negative values indicating a decrease with age.

### No evidence of a relationship between intelligence quotient with GWPC nor cortical thickness

We did not observe a significant influence of the total IQ score on either GWPC or cortical thickness, or on GWPC after correction on cortical thickness. This result was the same independently of the inclusion or exclusion of NYU, the multiple tests correction method, and was also observed in the EU-AIMS dataset.

### Grey-to-white percent contrast varies strongly across scanning sites

We observed a strong effect of acquisition centre on GWPC measured at 30% depth, capturing on average 69.8% of the variance across the brain (epsilon squared) in the ABIDE sample. In comparison, the age effect captured 0.567% of the variance, and the sex effect captured 0.027% of the variance. The full model captured on average R^2^=87.7% of the GWPC variance between individuals.

## DISCUSSION

We aimed at replicating recent reports of a difference in GWPC using a large number of individuals with a diagnosis of ASD and a non-ASD control group from the ABIDE project and from the EU-AIMS project. We did not observe evidence for such differences. We observed, however, that in a single ABIDE centre – NYU – the ASD and control groups had a strong, statistically significant difference in GWPC and cortical thickness. Our current interpretation is that this difference is most likely an artefact, although we have been unable to determine its nature. This issue is concerning, as NYU is one of the largest sites of ABIDE. The NYU data is in general of a very good quality, and the phenotype of the population does not differ significantly from that of other centres.

We have tried several different approaches to find what may be the cause of the NYU artefact. One possibility could be that because of its good data quality and large sample size, the NYU sample is able to capture a signal that the remaining 25 ABIDE centres fail to capture. However, through the EU-AIMS project we were able to access a large dataset, with very good quality obtained through extensive harmonisation efforts. We did not observe any statistically significant difference, neither in GWPC nor in cortical thickness. However, we did replicate in EU-AIMS the findings related to age and sex. We performed a series of exploratory analyses to try to better understand this artefact. These analyses did not change our main conclusion, and we have thus decided not to include them in the main result section but only as elements for discussion (the code we used is, however, available together with the code for the main results in the accompanying GitHub repository to facilitate replication).

Following the idea that NYU may be actually revealing a signal unavailable to the other sites, we aimed at comparing the surface maps of GWPC differences using a spin test (Alexander-Bloch et al. 2018). This test did not show a statistically significant correlation between the diagnosis effects of the two analyses (r=0.167, 95% CI [-0.027, 0.360], two-tailed p=0.092). We also tried in addition to the standard spin test to use a permutation test, shuffling the labels of the subjects of the ABIDE dataset (210 iterations), a method which is more statistically powered than the spin-test but requires more computation. This time, the correlation between diagnosis effect maps of ABIDE was statistically significant (95% CI [0.019, 0.314], two-tailed p = 0.0265), suggesting that a small effect may exist but not large enough and too diluted to be statistically significant on a specific region. It is not our aim, however, to keep trying any possible analyses until the results conform to our expectations, and these results are only meant to provide elements for further exploration.

A previous study of GWPC by Andrews et al. (2017) revealed regions of statistically significant decreases of GWPC in the ASD group. A more recent study by Olafson et al. (2021) reported, on the contrary, increases. The dataset analysed included a combination of open data along with private datasets. Their method is slightly different from the one used by Andrews et al. (2017) and by ourselves. Instead of computing GWPC, Olafson et al. (2021) fit a sigmoidal function to the grey-to-white interface, and use the slope of the sigmoidal (which they call the boundary contrast coefficient, BSC) to compare between groups: a steeper slope should correspond to a higher GWPC. We replicated their method by extracting surface intensities using FreeSurfer on a range of depth percentages going from -25% to 50% with incrementations of 6.25%. We used the R function nls for fitting the sigmoidal function and computing the BSC. Although the function still reported failure to convergence for some vertices, closer inspection showed that the fits seemed nevertheless acceptable. We thus included, as Olafson et al. (2021), all vertices. BSC maps correlated with GWPC and cortical thickness maps. We did not observe statistically significant BSC differences related to diagnosis after FDR correction neither in the ABIDE dataset (including or excluding NYU) nor in EU-AIMS. Using RFT correction, we found statistically significant clusters of diagnosis-related differences only on the NYU dataset. Overall, BSC led to similar results as GWPC, however, BSC analyses appeared to be less sensitive: they only revealed statistically significant differences for NYU, but not when NYU was included within the rest of ABIDE. In the analysis of their own dataset Olafson et al (2021) report an even stronger effect than what we found for NYU.

We cannot at this point conclude about the presence of differences in GWPC between individuals with ASD and non-ASD controls, we did not find any evidence for them. There are still many methodological decisions which can make results appear, or not, go in one direction or the other, especially when sample sizes are small. At this stage, using openly accessible data and making analysis scripts available is essential to better understand each other’s results. This is certainly not possible for all types of neuroimaging studies, however, those relying only on structural MRI and resting state functional MRI should use ABIDE systematically as a gold standard for replication. Through the combined effort of the research community, we should be able to understand the large dataset made available by the ABIDE project much more efficiently, which should in turn help us understand our own data. In particular, the group effect we observed in the NYU centre, which is likely an artefact, deserved further inspection and discussion. Our advice would be to systematically perform analyses using the ABIDE dataset including and excluding the NYU centre. All our code is available on GitHub: https://github.com/neuroanatomy/GWPC.

## ACKNOWLEDGMENTS

We thank the Fondation pour la Recherche Médicale, Grant number DEA20170637628, for financial support. This work uses EU-AIMS data obtained through project CONT.

## Supplemental Figures and Tables

### 1. Supplemental Figures

**Supplemental Figure 1.**
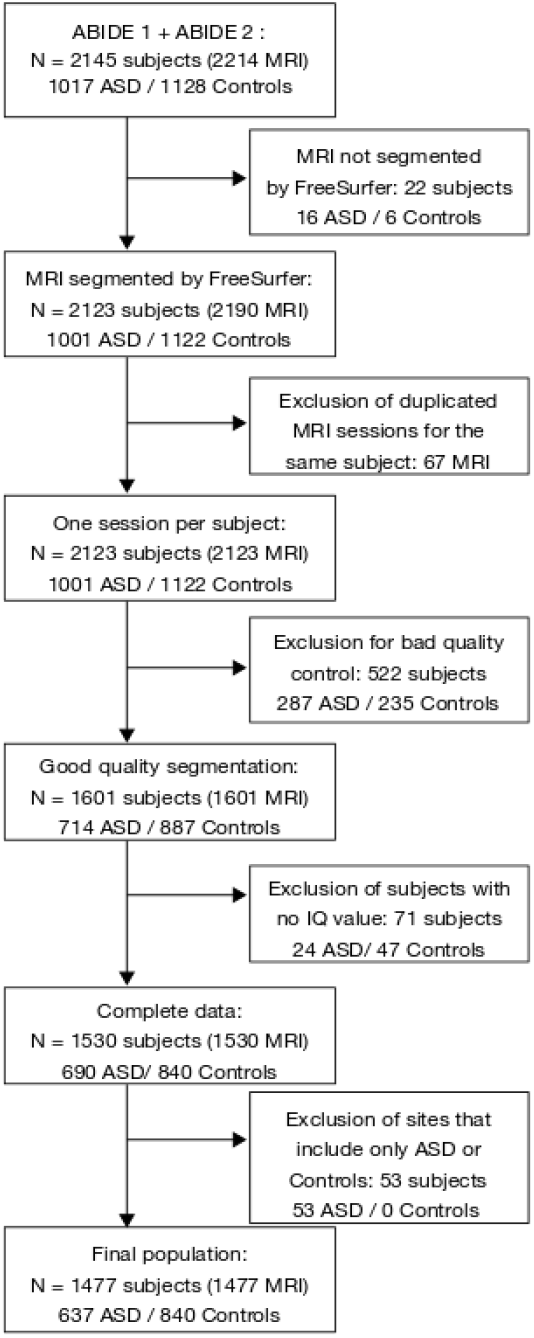
Flowchart describing the selection of the data to be analysed from the ABIDE 1 and 2 databases.

**Supplemental Figure 2.**
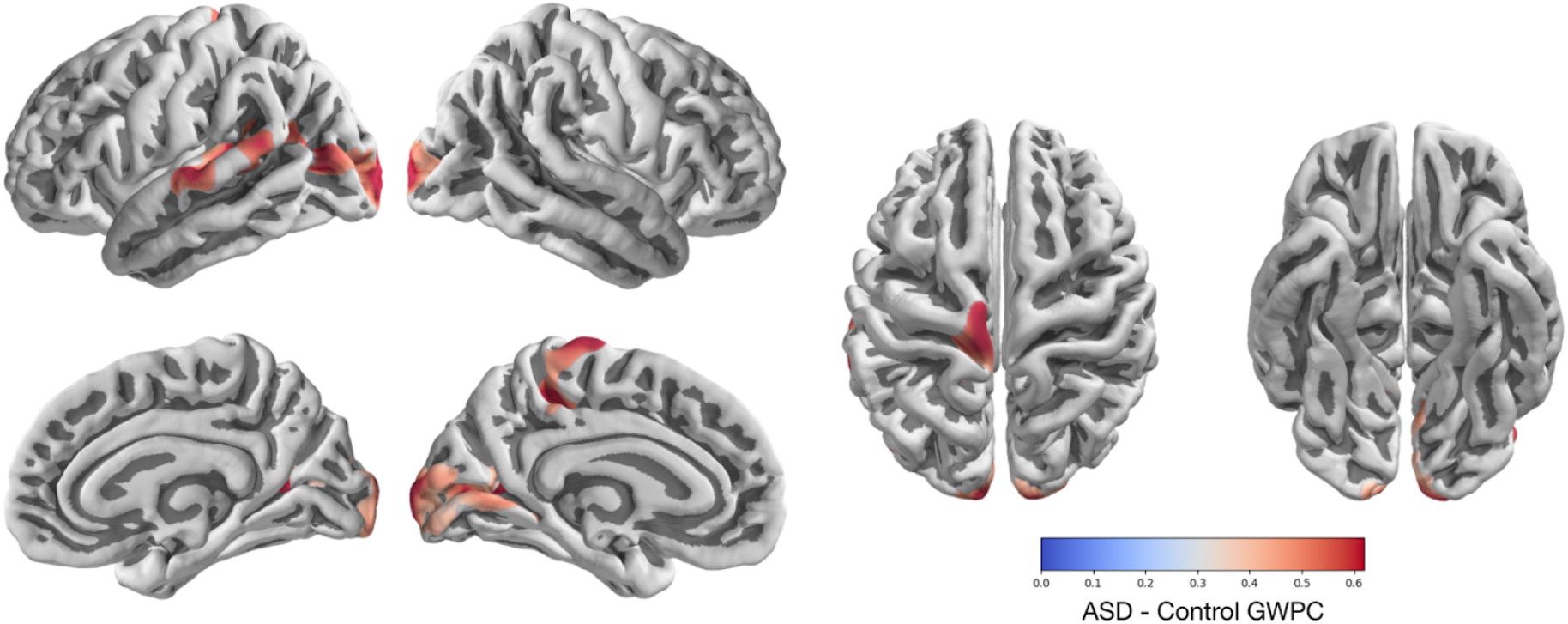
GWPC versus diagnosis, NYU site included. Statistically significant differences, corrected for multiple comparisons (RFT correction), were observed in several regions, including visual, auditory and somatosensory cortices. Positive values indicating increased contrast in ASD compared with controls. All differences were due to the inclusion of the NYU site, and were not significant when the site was excluded (see main manuscript).

**Supplemental Figure 3.**
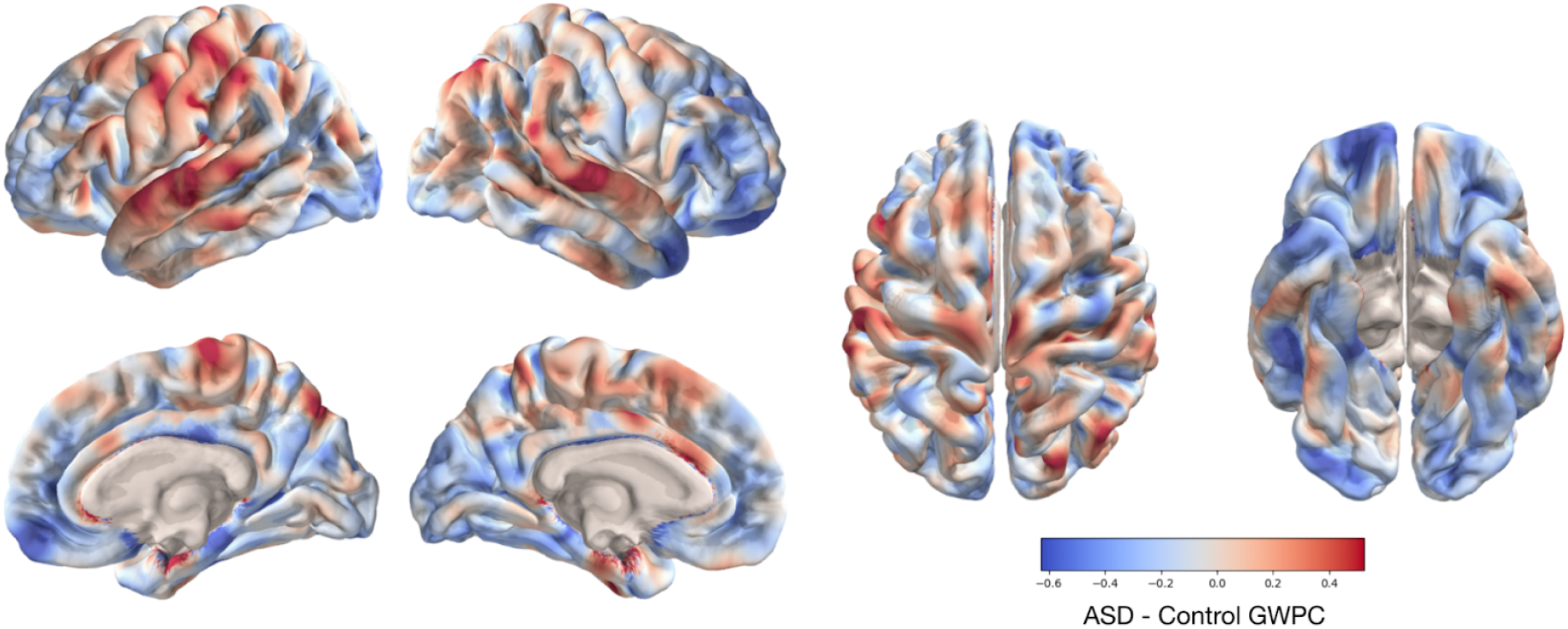
Non-thresholded GWPC versus diagnosis in the EU-AIMS dataset. GWPC measured at 30% GM depth. Red colour indicates larger contrast in subjects diagnosed ASD. The differences were not statistically significant.

**Supplemental Figure 4.**
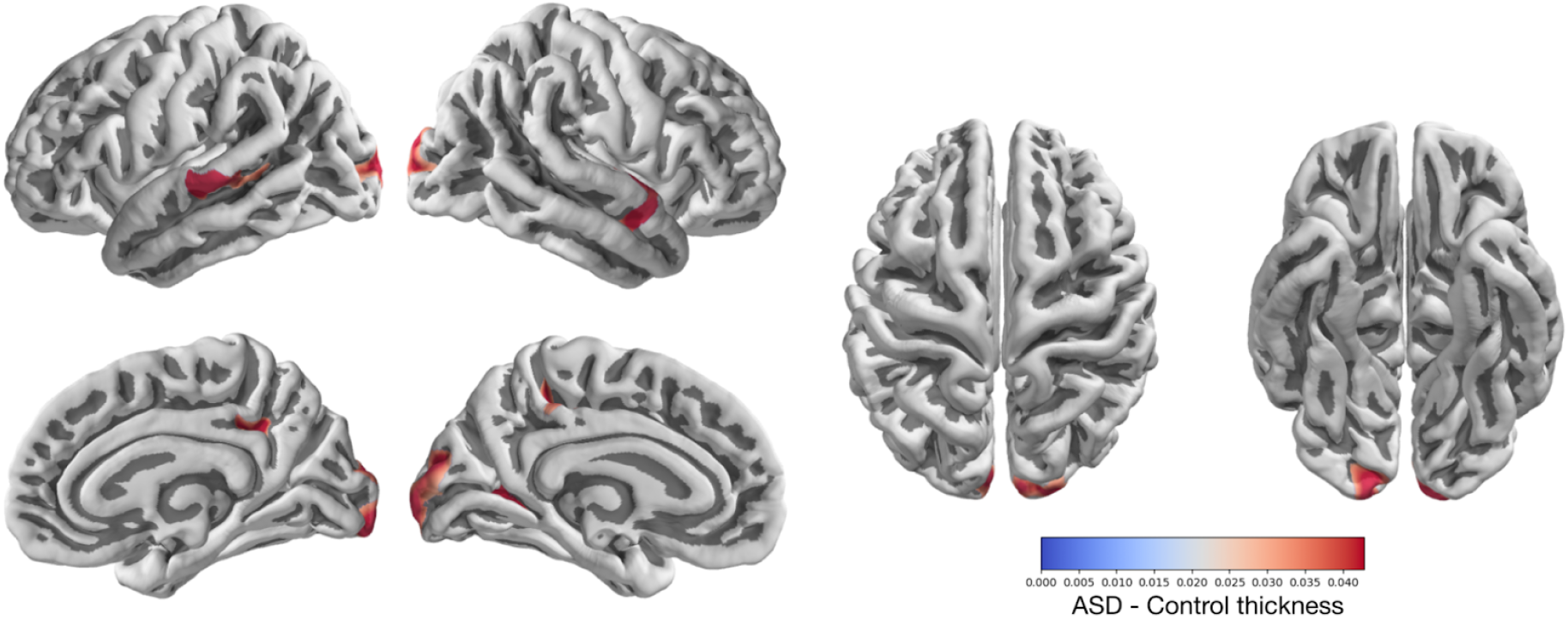
Cortical thickness versus diagnosis, NYU site included. Significantly thicker cortical regions were observed in the occipital and temporal lobes of subjects with ASD diagnosis. Only statistically significant results after correction for multiple comparisons are shown. Values are in millimetres, positive values: ASD>Control. No statistically significant differences were present when the NYU site was excluded.

**Supplemental Figure 5.**
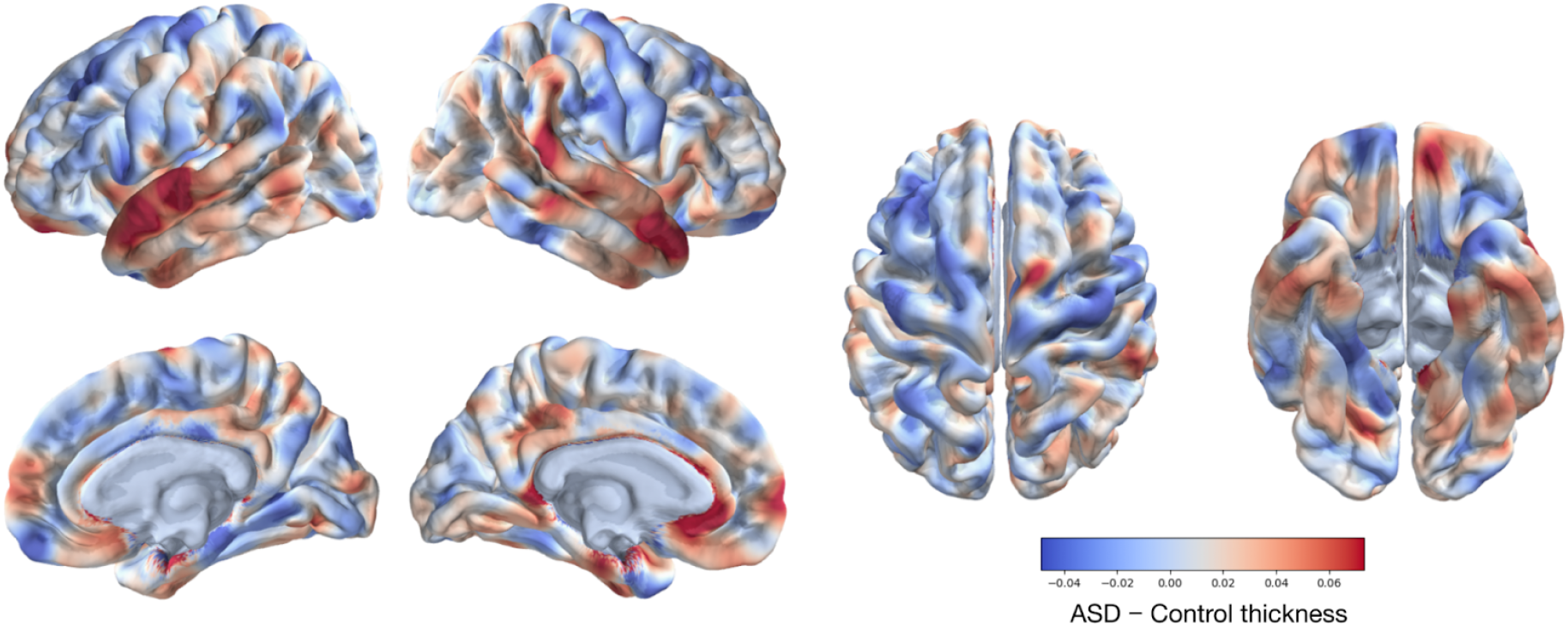
Non-thresholded cortical thickness versus diagnosis in the EU-AIMS dataset. Positive values indicate greater thickness in the ASD group. No differences were statistically significant.

**Supplemental Figure 6.**
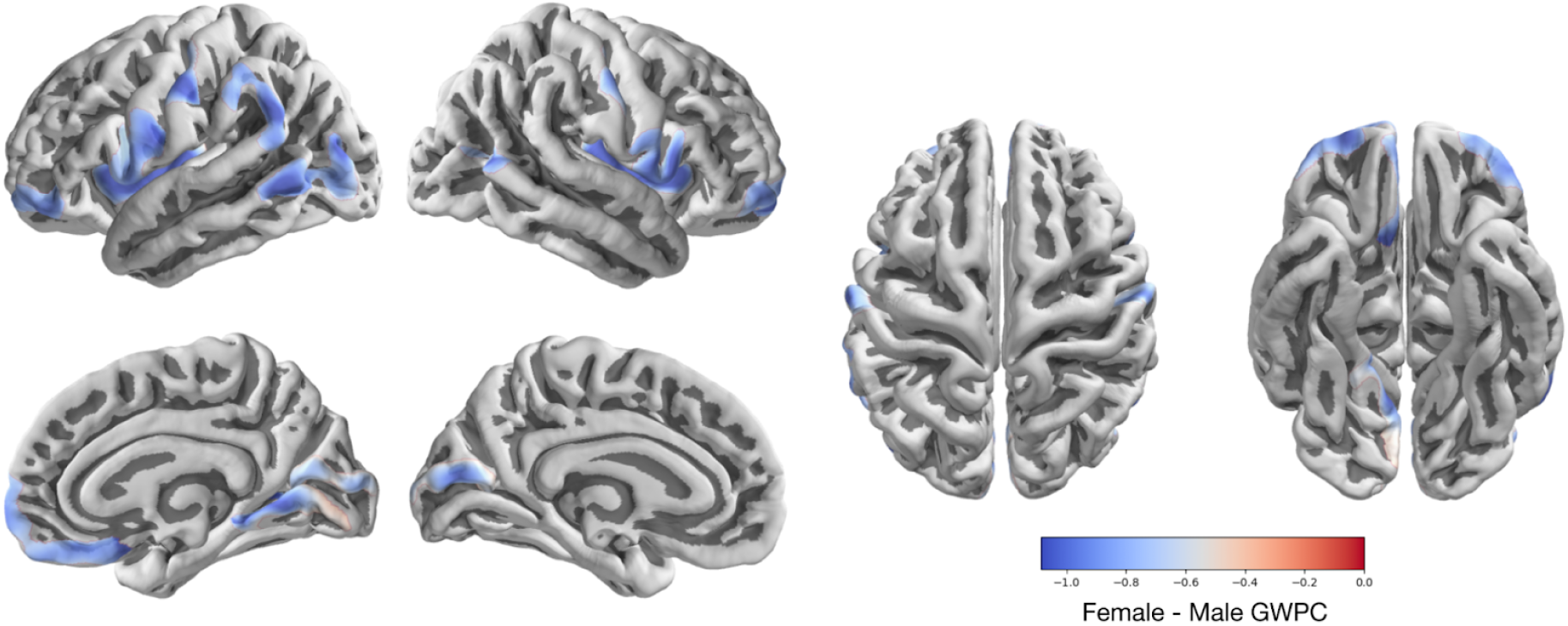
GWPC versus sex with cortical thickness as covariate, NYU site excluded. Only the significant results at 5% after correction for multiple tests by Monte Carlo are represented. Blue colour indicates regions with greater cortical thickness in females.

**Supplemental Figure 7.**
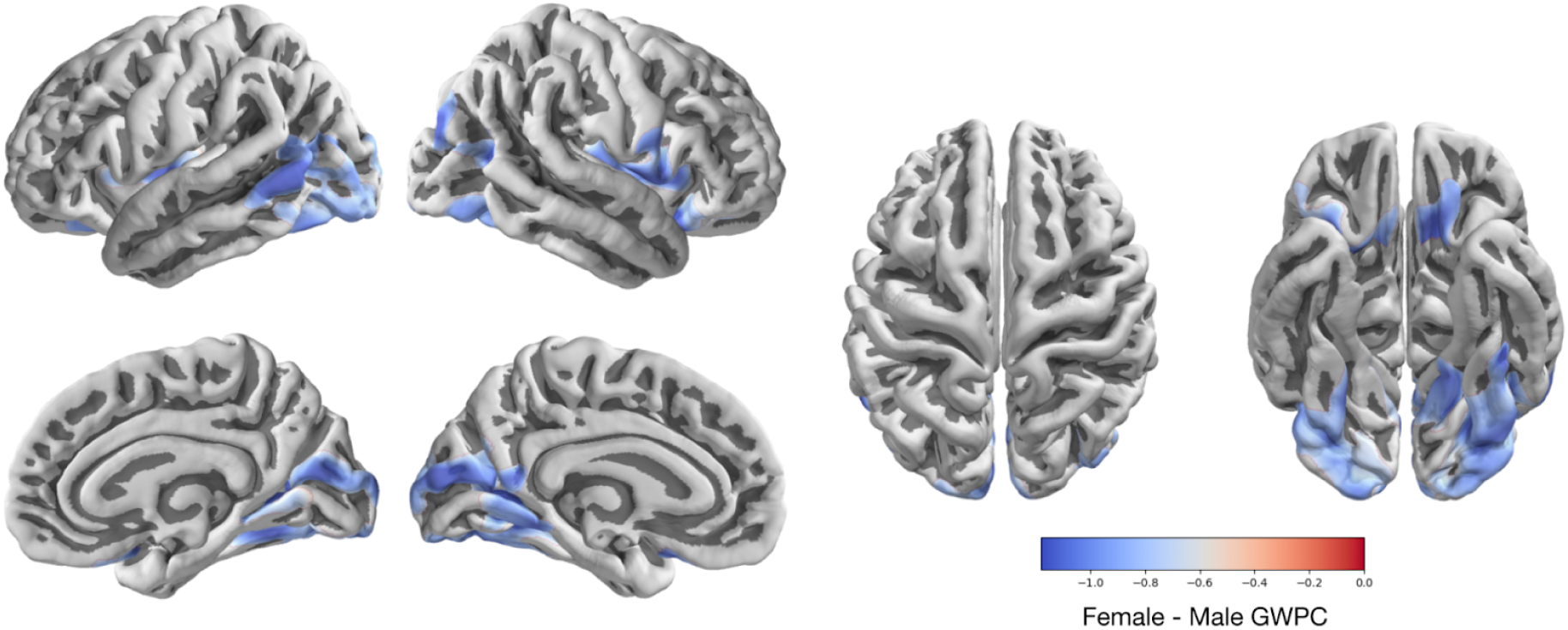
Significant GWPC versus sex in the EU-AIMS dataset. Correction for multiple comparisons using a RFT threshold. Blue colour indicates regions with larger contrast in females.

**Supplemental Figure 8.**
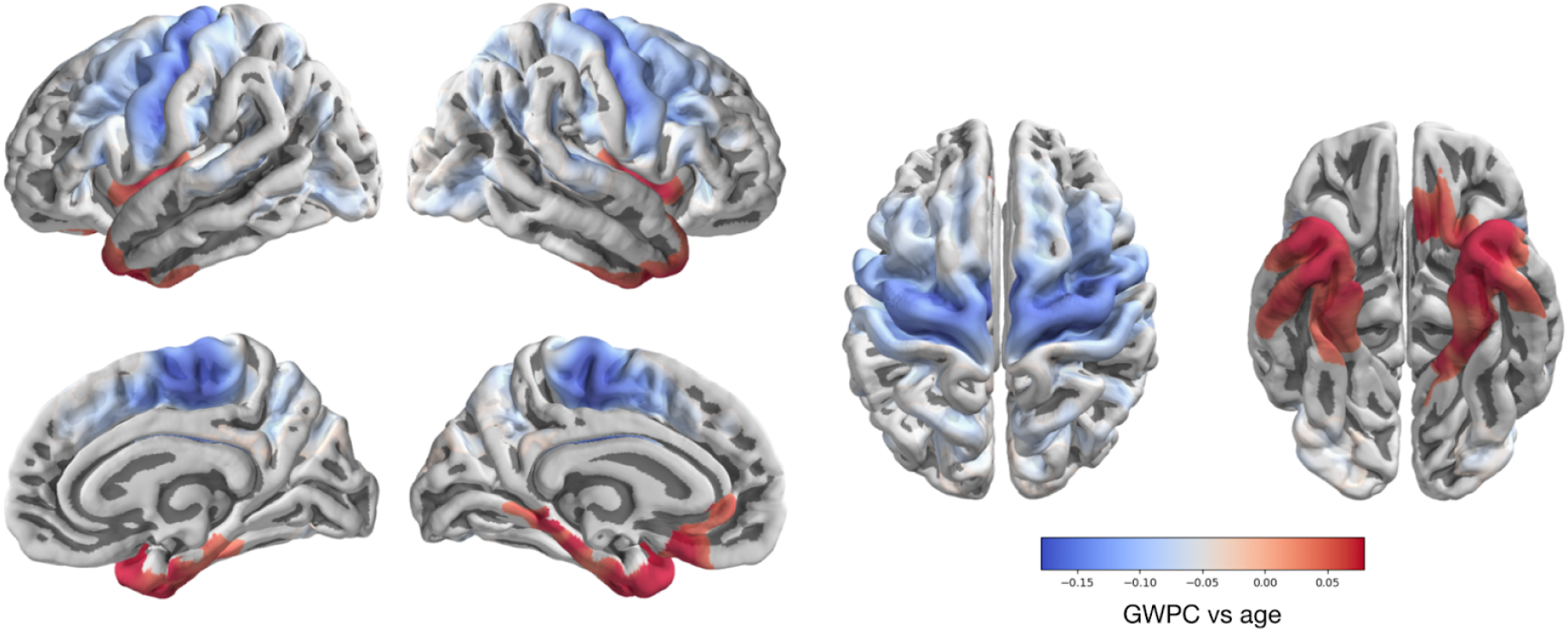
GWPC versus age, with cortical thickness as a covariate, NYU site excluded. Only significant results at 5% after correction by Monte Carlo for multiple tests are shown). From left to right: left hemisphere lateral and medial view, right hemisphere lateral and medial view. Note that the colour scale used is similar to that of GWPC difference maps with increasing age without inclusion of the cortical thickness, in order to facilitate visual comparison.

**Supplemental Figure 9.**
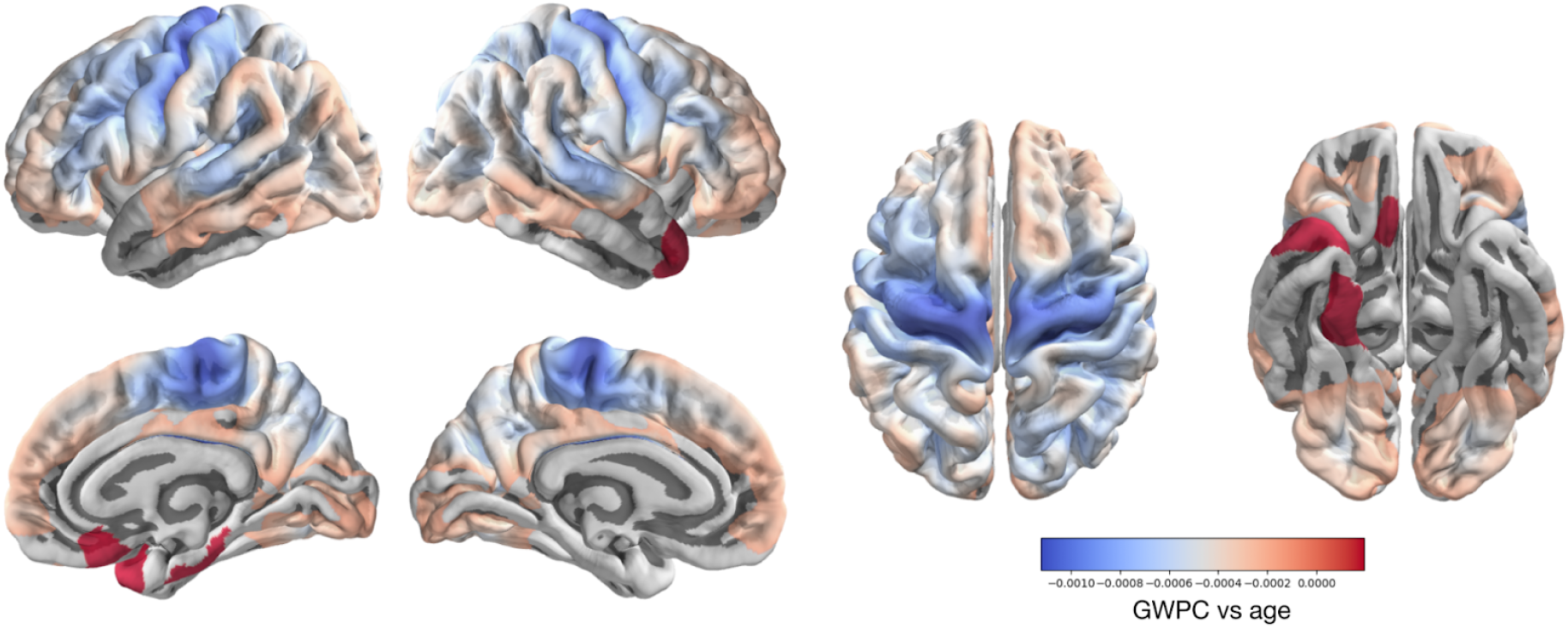
GWPC versus age in the EU-AIMS dataset. Correction for multiple comparisons using an RFT threshold. Red colour indicates regions where contrast increases with age.

### 2. Supplemental Tables

**Supplemental Table 1.** Distribution of Patients and Controls by Centre. 8 centres participated in Abide 1 and Abide 2 data: in bold in the table. Abbreviations corresponding to the centres: BNI_1 (Barrow Neurological Institute), ETH_1 (ETH Zurich), GU (Georgetown University), IP_1 (Institut Pasteur and Robert Debre Hospital), IU_1 (Indiana University), KKI_1 (Kennedy Krieger Institute), NYU (NYU) Langone Medical Center), OHSU (Oregon Health and Science University), OLIN_1 and OLIN_2 (ONRC: Olin Neuropsychiatry Research Center, Institute of Living at Hartford Hospital, samples 1 and 2), SDSU (San Diego State University), TRINITY (Trinity Center for Health Sciences), UCD_1 (University of California Davis), UCLA_1 (University of California Los Angeles, sample 1), UCLA_2 (University of California Los Angeles, sample 2), USM (University of Utah, School of Medicine), CALTECH (California Institute of Technology), CMU (Carnegie Mellon University), LEUVEN_1 (University of Leuven, sample 1), LEUVEN_2 (University of Leuven, sample 2), MAX_MUN (Ludwig Maximilians University Munich), PITT (University of Pittsburgh School of Medicine), UM (University of Michigan), Yale (Yale Child Study Center).

| Site | ASD<br>(N = 636 [86 ♀, 550 ♂]) | Controls<br>(n = 839 [202 ♀, 637 ♂]) |
| --- | --- | --- |
| BNI_1 (n = 58, [58 ♂]) | N= 29 [29 ♂] | N = 29 [29 ♂] |
| ETH_1 (n = 31, [31 ♂]) | N = 9, [9 ♂] | N = 22, [22 ♂] |
| GU_1 (n = 78, [50 ♂, 28 ♀]) | N = 32 [27 ♂, 5 ♀] | N = 46 [23 ♂, 23 ♀] |
| IP_1 (n = 6, [4 ♂, 2 ♀]) | N = 4 [3 ♂, 1 ♀] | N = 2 [1 ♂, 1 ♀] |
| IU_1 (n = 7 [5 ♂, 2 ♀]) | N = 1 [1 ♀] | N = 6 [5 ♂, 1 ♀] |
| KKI_1 (n = 122 [79 ♂, 43 ♀]) | N= 32 [24 ♂, 8 ♀] | N = 90 [55 ♂, 35 ♀] |
| KKI_2 (n = 49 [35 ♂, 14 ♀]) | N = 12 [8 ♂, 4 ♀] | N = 37 [27 ♂, 10 ♀] |
| KKI_3 (n = 57 [39 ♂, 18 ♀]) | N = 15 [12 ♂, 3 ♀] | N = 42 [27 ♂, 15 ♀] |
| NYU (n = 227 [186 ♂, 41 ♀]) | N= 101 [86 ♂, 15 ♀] | N = 126 [100 ♂, 26 ♀] |
| OHSU (n = 117 [82 ♂, 35 ♀]) | N = 47 [41 ♂, 6 ♀] | N = 70 [41 ♂, 29 ♀] |
| OLIN_1 (n = 20 [17 ♂, 3 ♀]) | N = 9 [8 ♂, 1 ♀] | N= 11 [9 ♂, 2 ♀] |
| ONRC_2 :OLIN 2<sup>nd</sup> sample (n = 48 [32 ♂, 16 ♀]) | N = 17 [14 ♂, 3 ♀] | N = 31 [18 ♂, 13 ♀] |
| SDSU (n = 67 [55 ♂, 12 ♀]) | N = 32 [25 ♂, 7 ♀] | N = 35 [30 ♂, 5 ♀] |
| TRINITY (n = 47 [47 ♂]) | N = 24 [24 ♂] | N = 23 [23 ♂] |
| TCD_1 (TRINITY 2<sup>nd</sup> sample) (n = 36 [36 ♂]) | N = 16 [16 ♂] | N = 20 [♂, ♀] |
| UCD_1 (n = 27 [20 ♂, 7 ♀]) | N = 15 [12 ♂, 3 ♀] | N = 12 [8 ♂, 4 ♀] |
| UCLA Abide1 (n = 83 [71 ♂, 12 ♀]) | N = 46 [40 ♂, 6 ♀] | N = 37 [31 ♂, 6 ♀] |
| UCLA Abide2 (n = 19 [15 ♂, 4 ♀]) | N = 9 [9 ♂] | N = 10 [6 ♂, 4 ♀] |
| USM (n = 125 [120 ♂, 5 ♀]) | N = 67 [65 ♂, 2 ♀] | N = 58 [55 ♂, 3 ♀] |
| CALTECH (n = 34 [27 ♂, 7 ♀]) | N = 15 [12 ♂, 3 ♀] | N = 19 [15 ♂, 4 ♀] |
| CMU (n = 22 [16 ♂, 6 ♀]) | N = 11 [8 ♂, 3 ♀] | N = 11 [8 ♂, 3 ♀] |
| LEUVEN (n = 58 [51 ♂, 7 ♀]) | N = 28 [26 ♂, 2 ♀] | N = 30 [25 ♂, 5 ♀] |
| MAX_MUN (n = 35 [28 ♂, 7 ♀]) | N = 14 [11 ♂, 3 ♀] | N = 21 [17 ♂, 4 ♀] |
| PITT (n = 54 [46 ♂, 8 ♀]) | N = 27 [23 ♂, 4 ♀] | N = 27 [23 ♂, 4 ♀] |
| UM (n = 24 [20 ♂, 4 ♀]) | N = 11 [8 ♂, 3 ♀] | N = 13 [12 ♂, 1 ♀] |
| YALE (n = 24 [17 ♂, 7 ♀]) | N = 13 [10 ♂, 3 ♀] | N = 11 [7 ♂, 4 ♀] |

**Supplemental Table 2.**
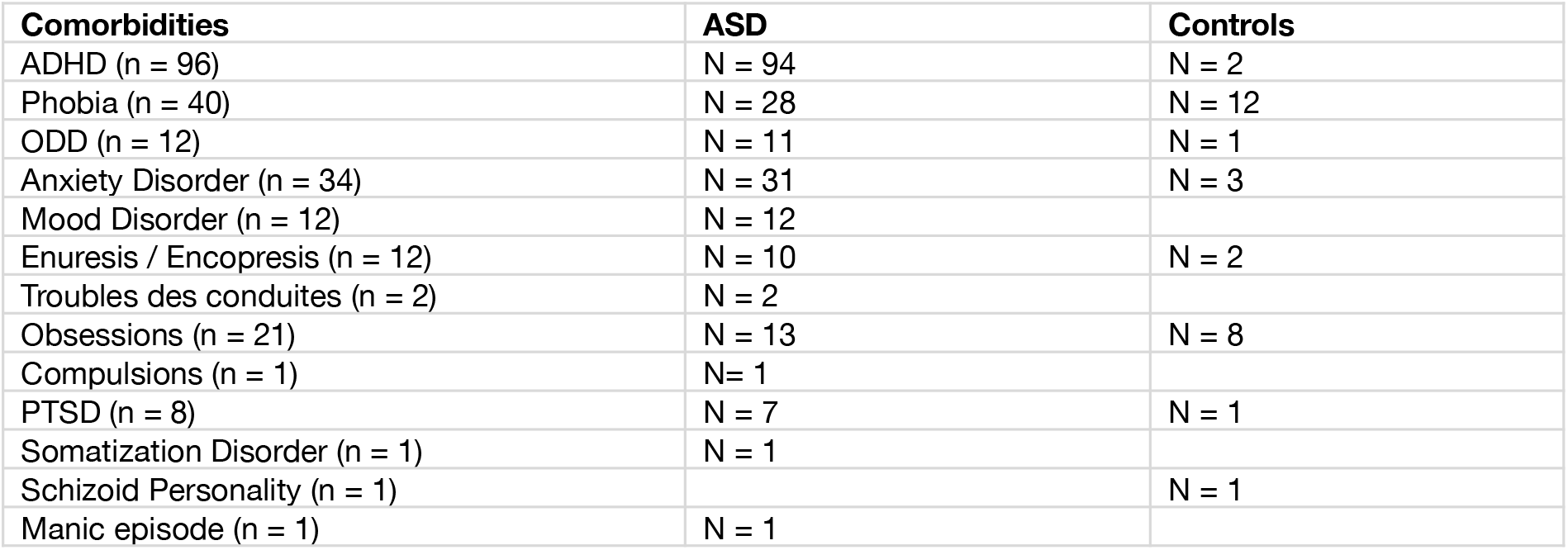
Summary of comorbidities identified in the studied population. ADHD = Attention Deficit with Hyperactivity Disorder. The phobia item includes patients suffering from social phobia and specific phobia. ODD: Oppositional Defiant Disorder. The “anxiety disorder” item includes patients with anxiety, separation anxiety, and generalized anxiety disorder. The 3 identified controls had a generalised anxiety disorder. Regarding the Enuresis / Encopresis item: 9 individuals had enuresis (7 ASD patients and 2 controls), and 3 ASD patients had encopresis. PTSD = Post Traumatic Stress Disorder. No comorbidities of schizophrenia, antisocial personality disorder, anorexia nervosa, bulimia, or substance abuse were found in either controls or ASD subjects.

**Supplemental Table 3.**
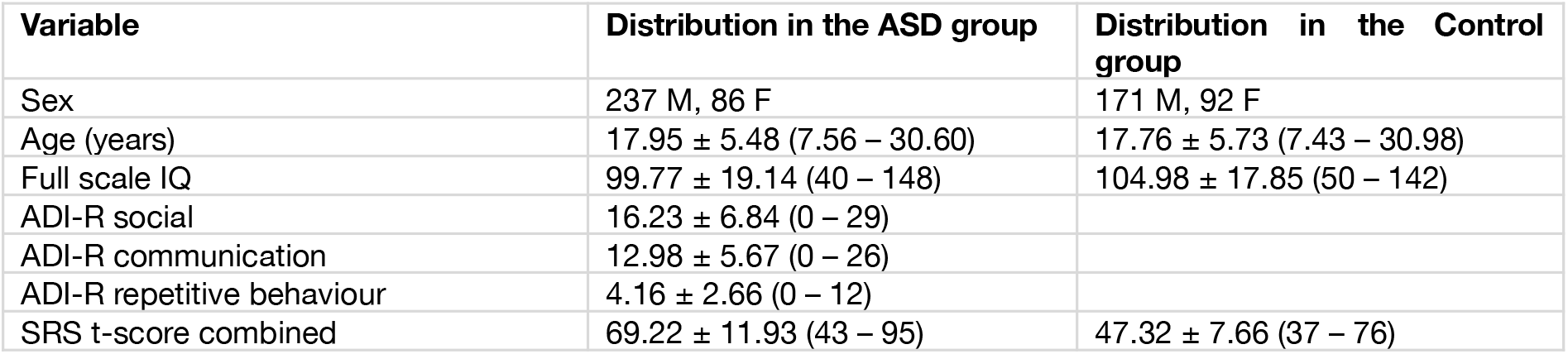
Characteristics of the study sample for the EU-AIMS dataset. Results expressed as mean ± standard deviation (minimum, maximum). ADI-R: Autism Diagnostic Interview-Revised, SRS: Social Responsiveness Scale. ADI-R scores are missing for 16 ASD subjects, SRS t-score is missing for 162 ASD subjects and 49 typical controls.

**Supplemental Table 4.** Distribution of Patients and Controls by Centre for the EU-AIMS dataset. Abbreviations corresponding to the centres: Cambridge (University of Cambridge), KCL (King’s College London), Mannheim (Central Institute of Mental Health Mannheim), Nijmegen (Radboud University Nijmegen Medical Centre), Rome (Università Campus Bio-Medico di Roma), Utrecht (University Medical Centre Utrecht).

| Site | ASD<br>(N = 323 [86 ♀, 237 ♂]) | Controls<br>(n = 263 [92 ♀, 171 ♂]) |
| --- | --- | --- |
| Cambridge (n = 74 [49 ♂, 25 ♀]) | N = 44 [28 ♂, 16 ♀] | N = 30 [21 ♂, 9 ♀] |
| KCL (n = 173 [121 ♂, 52 ♀]) | N = 100 [80 ♂, 20 ♀] | N = 73 [41 ♂, 32 ♀] |
| Mannheim (n = 63 [46 ♂, 17 ♀]) | N = 29 [22 ♂, 7 ♀] | N = 34 [24 ♂, 10 ♀] |
| Nijmegen (n = 164 [114 ♂, 50 ♀]) | N = 95 [68 ♂, 27 ♀] | N = 69 [46 ♂, 23 ♀] |
| Rome (n = 36 [26 ♂, 10 ♀]) | N = 19 [14 ♂, 5 ♀] | N = 17 [12 ♂, 5 ♀] |
| Utrecht (n = 76 [52 ♂, 24 ♀]) | N = 36 [25 ♂, 11 ♀] | N = 40 [27 ♂, 13 ♀] |

**Supplemental Table 5.** Clusters with statistically significant decrease in GWPC among females at 30% depth of GM. The number of resolution elements corresponds to the number of independent observations within the cluster. P cluster is the p corrected value of the cluster.

| Cortical regions | Hemisp here | Number of vertex points | Number of resolution elements | Cluster corrected p-value | Mean difference (%) |
| --- | --- | --- | --- | --- | --- |
| insula, pre- and postcentral gyri, lateral orbitofrontal cortex, pars orbitalis and opercularis of the inferior frontal gyrus, supramarginal gyrus, superior temporal gyrus | Left | 8378 | 15.108 | <1e-100 | -0.912 |
| Lateral occipital cortex, middle and lower temporal gyrus, cuneus, pericalcarine region, lower parietal cortex | Left | 4841 | 10.52 | 2.35e-14 | -0.768 |
| Medial and lateral orbitofrontal cortex, rostral portion of the middle frontal gyrus, pars orbitalis of the inferior frontal gyrus, frontal pole, superior frontal cortex, pars triangularis of the inferior frontal gyrus | Left | 3087 | 6.142 | 4.2e-08 | -0.809 |
| Lingual cortex, isthmus of the cingulate cortex, precuneus, parahippocampal gyrus | Left | 1900 | 4.932 | 2.24e-06 | -0.819 |
| Supramarginal gyrus, superior temporal gyrus | Left | 1380 | 3.199 | 0.000666 | -0.797 |
| precuneus, superior parietal gyrus | Left | 1152 | 2.272 | 0.014 | -0.753 |
| Fusiform cortex | Left | 966 | 2.258 | 0.0146 | -0.679 |
| lingual, cuneus, pericalcarine, isthmus of the cingulate cortex, parahippocampal, precuneus, lateral occipital | Right | 3589 | 10.423 | 3.24e-14 | -0.815 |
| insula, precentral, pars opercularis, postcentral, superior temporal, lateral orbitofrontal | Right | 5648 | 10.036 | 1.16e-13 | -0.98 |
| middle temporal, inferior parietal, inferior temporal, lateral occipital, Banks of the Superior Temporal Sulcus | Right | 2133 | 4.752 | 4.03e-06 | -0.863 |
| caudal anterior cingulate, posterior cingulate, rostral anterior cingulate | Right | 1411 | 4.746 | 4.11e-06 | -0.776 |
| rostral middle frontal, frontal pole, superior frontal, pars orbitalis, medial orbitofrontal | Right | 2498 | 4.568 | 7.39e-06 | -0.802 |
| fusiform, inferior temporal | Right | 1387 | 3.572 | 0.000195 | -0.773 |
| medial orbitofrontal | Right | 864 | 3.564 | 0.0002 | -0.921 |

**Supplemental Table 6.** Clusters with significant decrease in GWPC with age at 30% depth of GM. The number of resolution elements corresponds to the number of independent observations within the cluster. P cluster is the p corrected value of the cluster.

| Cortical regions | Hemisphere | Number of vertex points | Number of resolution éléments | Cluster corrected p-value | Mean difference (%) |
| --- | --- | --- | --- | --- | --- |
| Upper and middle frontal cortex, pre- and postcentral gyri, superior and inferior parietal cortex, supramarginal gyrus, precuneus, rostral middle frontal cortex, lateral occipital cortex, superior, middle, inferior and transverse temporal gyri, lingual cortex, paracentral sulcus, pars opercularis, fusiform cortex, posterior and anterior cingulate cortex, lateral and medial orbitofrontal cortex, superior temporal sulcus, isthmus of cingulate cortex, pars triangularis, insula, pericalcarine region, cuneus, pars orbitalis, frontal pole | Left | 130433 | 306.637 | <1e-100 | -0.133 |
| Temporal pole, entorhinal cortex, fusiform cortex, upper and lower temporal gyri | Left | 1675 | 14.561 | <1e-100 | 0.067 |
| Insula | Left | 1262 | 2.418 | 0.00865 | 0.066 |
| Upper and middle caudal frontal cortex, pre- and postcentral gyri, paracentral sulcus, superior and inferior parietal cortex, supramarginal gyrus, precuneus, middle frontal cortex, latero-occipital cortex, superior, middle, inferior and transverse temporal gyri, lingual cortex, lateral and medial orbitofrontal cortex, posterior and anterior cingulate cortex, fusiform gyrus, pars opercularis, pars triangularis, superior temporal sulcus, isthmus of the cingulate cortex, pericalcarine region, cuneus, insula, pars orbitalis, frontal pole, parahippocampal cortex | Right | 130533 | 302.551 | <1e-100 | -0.134 |
| Temporal pole, parahippocampal cortex, entorhinal cortex, fusiform gyrus, upper and middle temporal gyri, insula | Right | 2491 | 14.452 | <1e-100 | 0.055 |
| Insula | Right | 1428 | 2.424 | 0.00846 | 0.066 |
| Medial orbito-frontal cortex | Right | 319 | 2.247 | 0.0151 | 0.064 |

